# Analysis of insertion mutants in distal α9 portion of C-terminal heptad repeat (CHR) of HIV-1 gp41 subunit

**DOI:** 10.1101/2020.03.30.017129

**Authors:** Hongyun Wang, Jiping Song, Yasushi Kawaguchi, Jun-ichiro Inoue, Zene Matsuda

## Abstract

We have made insertion mutants in α9 of HXB2 gp41 and observed similar phenotypes like recent JRFL mutants: insertion of alanine (653+A), but not glutamine (653+Q), severely attenuated membrane fusion. To understand the underlying mechanism, we performed the fusion inhibition assay by corresponding mutant C34 peptides. Both mutant C34 peptides added at the beginning of the coculture of the effector and target cells showed less efficient inhibition of membrane fusion, which was similar to wildtype C34 added after 30 min of coculture, indicating slow association of mutant C34 peptides with the N-terminal heptad region of gp41. Due to uninterpretable CD profiles of C34 and N36, we tested the longer peptide pairs (N46 and C42) and observed CD profiles indicative of weak α-helix formation. The melting temperatures for N46-C42 pairs of 653+A, 653+Q, and wild type were 56.8 °C, 59.8°C, and 96°C, respectively. Taken together, our data suggested that the phenotypic difference in membrane fusion between 653+A and 653+Q (or wild type) was not based on the stability of the six-helix bundle (6HB), but due to differences in the kinetics of 6HB formation. Further, we examined additional insertions (E, R, I, and L) at position 653, for which only I and L showed fusion recovery similar to Q, suggesting that the polar nature of glutamine was not a phenotypic determinant.

## Introduction

Membrane fusion is the first step of the HIV-1 life cycle, mediated by the viral envelope glycoprotein (Env) (1,2). Env is first synthesized as a heavily glycosylated precursor protein and cleaved into the mature surface envelope glycoprotein (gp120) and transmembrane envelope glycoprotein (gp41). In its native state, trimeric gp41 non-covalently associates with a gp120 trimer (3,4).

According to the current model, attachment of the gp120 subunit to the cell surface receptor/coreceptor (CD4 and CXCR4/CCR5) complex initiates conformational changes in gp120/gp41 needed for membrane fusion. The gp41 subunit is a member of the class I fusion proteins, which undergoes drastic conformational changes during the membrane fusion process. The fusion peptide, located at the N-terminus of gp41, is inserted into the target membrane upon formation of the extended coiled-coil of the N-terminal heptad repeats (NHRs) of gp41. The C-terminal heptad repeats (CHRs) of gp41bends over in an anti-parallel fashion onto the grooves formed on the NHR coiled coil to create a six-helix bundle (6HB) structure. This packing interaction of NHR and CHR brings the viral and cellular membranes closer, leading to membrane fusion (5-7). Membrane fusion is believed to proceed through hemi-fusion, initial fusion pore formation, pore enlargement, and eventual merging of the virus and cell membranes (8-13). However, the structures of Env in the intermediate states and their relationship with lipid bilayers are still unknown. For example, the physical relationship between forming 6HB and lipids at different time points remains elusive.

In a previous study, we performed alanine insertion mutagenesis in the gp41 CHR region (14) and conducted syncytia formation or the dual split protein (DSP) assay, a more quantitative kinetic fusion assay, to analyze the ability of the mutants to mediate membrane fusion. The DSP assay employs a pair of split luciferase-based reporters (dual split protein), which has been widely used to analyze the membrane fusion of HIV-1 and other viruses (15-20). Previous results have shown that alanine insertion in the distal portion of CHR (beyond position 647) of HXB2 gp41subunit (where CHR leaves the four-helix collar formed by α6, α7, α8, and α9) was more tolerable than the preceding regions. Furthermore, fusion was improved when insertion points were moved towards the C-terminal of CHR. These data suggested that 6HB formation, the zipping between NHR and CHR, is likely initiated by interactions between the distal end of NHR and the proximal end of CHR, as previously proposed by Markosyan et al (21).

In another previous study, we extended mutagenesis into more downstream of CHR using JRFL Env, and found the most of downstream insertions were also tolerable, except for 644+A. Interestingly, mutation of the inserted alanine to glutamine (644+Q) recovered the fusion activity. We hypothesized that the polar nature of Q, which facilitated formation of the hydrogen bond network with other glutamine residues in 6HB, was important for efficient membrane fusion (22).

In this study, we generated corresponding insertion mutants in HXB2 gp41 (653+A or Q of HXB2 corresponding to 644+A or Q in JRFL, respectively) and examined their phenotypes. As JRFL and HXB2 have highly conserved sequences in this region, we observed similar phenotypes in the fusion profile. To analyze the mechanism underlying the difference in fusion phenotypes between 653+A and 653+Q, we used synthetic peptides corresponding to mutants and compared their membrane fusion inhibitory activity and biophysical properties. Our data suggested that the kinetics of 6HB formation, not 6HB stability, may be the determinant of membrane fusion. Further, our analyses of the insertion mutants that replaced the inserted Q residue with other amino acids (E, R, I, and L) suggested that the polar nature of Q may not be a critical factor for these phenotypes.

## Results

### Effect of insertion mutation in distal portion of α9 inHXB2 CHR on cell-cell fusion activity

Based on the previous study in JRFL (22), we generated corresponding alanine insertion mutants in the distal portion of α9 using HXB2Env (650+A to 655+A, indicated by arrowheads in Fig. 1A). We also constructed corresponding insertion mutants of 653+Q. As for the fusion assays, we used the syncytia formation assay or quantitative kinetic fusion assay called DSP assay. Similar to JRFL, 653+A in HXB2 showed severe fusion defects by both syncytia and DSP assays. Although attenuated in some mutants, membrane fusion activity was retained in all other alanine insertion mutants. Moreover, substitution of the inserted alanine to Q, 653+Q, recovered the fusion activity (Fig.1B and C). Immunoblotting results showed comparable expression and processing level of 653+A and 653+Q mutants to the wild-type protein (Fig.4). Taken together, the findings suggest that the fusion defect in HXB2 mutants occurred within the fusion process, similar to JRFL mutants.

**Figure 1.**
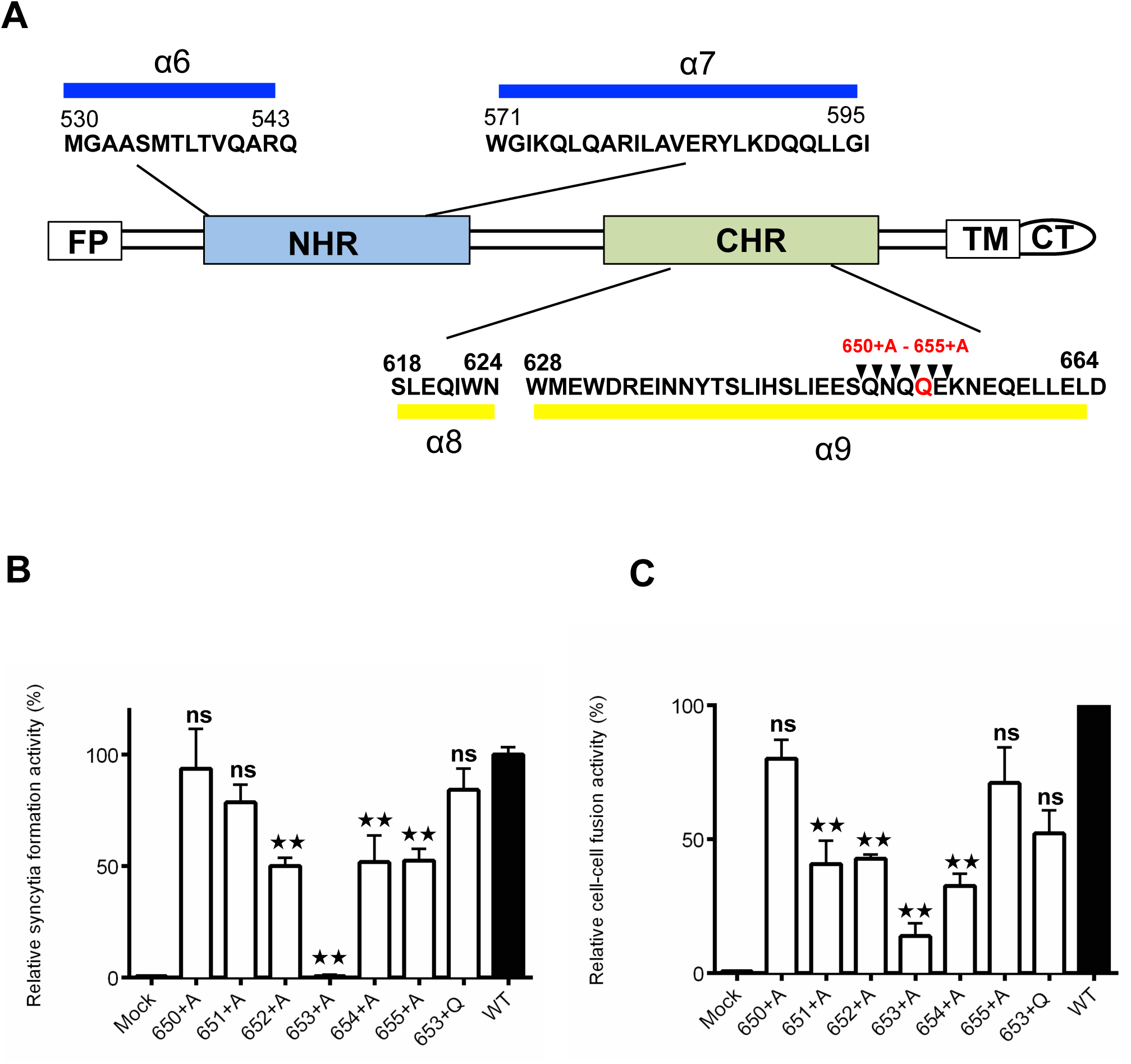
Analysis of alanine insertion mutants of gp41. (A) Schematic representation of HIV-1 gp41. FP, fusion peptide; NHR, N-terminal heptad repeat region; CHR, C-terminal heptad repeat region; TM, transmembrane domain; CT, cytoplasmic tail. The positions of α-helices are shown by colored boxes (blue and yellow) with the corresponding sequences. The insertion sites are indicated by arrowheads above the gp41 sequence. Mutants are named by the position of the inserted residue (such as 650+A). The numbering is based on HXB2 Env. The Q residue at the 653 position is shown in red. (B) Syncytia formation assay of the mutants. The expression vector for each mutant was transfected into 293CD4 cells, and syncytia formation was evaluated by microscopy. The relative degree of syncytia formation was shown by dividing the number of nuclei in the syncytia by the total number of nuclei in the observed fields. The wild type value was set as 100%. Error bars represent the standard deviation of results from three fields. Statistical significance was calculated by t test.**: p<0.01; ns indicates no significant differences vs. wild-type (WT) protein.(C) DSP assay at 2 h after coculture. Statistical analysis was performed using Prism 6 software. Error bars represent the standard deviation of triplicate experiments. **: p<0.01; ns indicates no significant difference vs. wild-type (WT) protein.

### Analysis of complex formation between N36 and wild type or mutant C34 peptides

CD spectra have been previously used to analyze the formation of α-helical 6HB between NHR and CHR, which is mechanistically and thermodynamically correlated with HIV-1 fusion (23-26). Because mutations that weaken the NHR and CHR interaction may affect fusion, we investigated the difference between 653+A and 653+Q using synthesized C34 peptides of these mutants. In contrast to wild-type C34, the mixture of N36 and mutant C34 peptides did not show two dips in the CD spectra, indicative of α-helix formation (Fig.2A), suggesting either a poor interaction between N36 and mutant C34 or poor helix formation. Moreover, thermal stability of the mixture was evaluated by monitoring the CD signal at 222 nm. However, no temperature shift for N36 plus C34 mutants was detected (Fig.2B).

**Figure 2.**
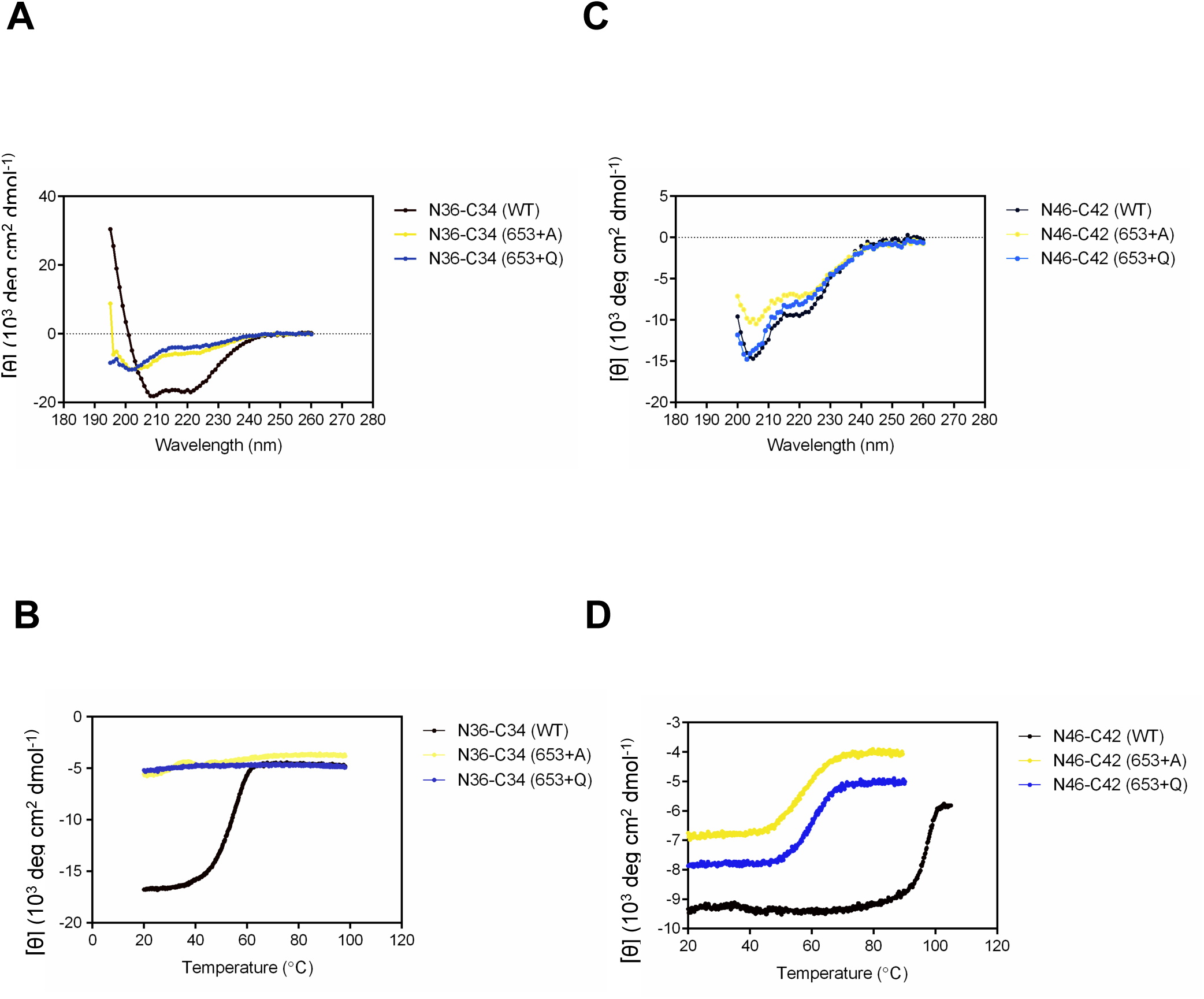
Biophysical properties of complexes formed between NHR-and CHR-derived peptides by CD spectrographic analysis. (A) CD spectra of complexes formed by N36 and C34 (black), N36 and C34 (653+A) (yellow), or N36 and C34 (653+Q)(blue) at a peptide concentration of 10 µM in PBS at room temperature. (B) The thermal stability of complexes formed by N36 with C34 (black) and its mutants (653+A, yellow; and 653+Q, blue) was evaluated by monitoring the CD signal at 222 nm.(C) CD spectra of complexes formed by N46 and C42 (black), N46 and C42 (653+A) (yellow), or N46 and C42 (653+Q)(blue) at a peptide concentration of 10 µM in PBS at room temperature. (D) The stability of complexes formed by N46 with C42 (black) and its mutants (C42 (653+A), yellow and C42(653+Q), blue) was measured at 222 nm by thermal denaturation analysis.

### MutantC34 peptides inhibited membrane fusion

To determine whether the mutant C34 peptides could interact with NHR, we examined the inhibitory effect of the mutant C34 peptides (653+Aand 653+Q) on membrane fusion by the DSP assay. In the DSP assay, measurement of split *Renilla* luciferase (RL) re-association upon membrane fusion (pore formation) served as a reporter of pore formation between co-cultured effector and target cells. Additionally, another mutant C34 peptide, 647+A, with an alanine insertion at position 647, upstream of position 653 was included (14). Further, wild-type or mutant C34 peptides (4μM) were added at the start or 30 min after co-culture. Results showed that when added at the start of the co-culture, wild type C34 completely inhibited membrane fusion. Both mutant C34 (653+A, or 653+Q) showed about 70% inhibition of membrane fusion, while C34 (647+A) showed only 30% inhibition (Fig.3A). When added 30 min after co-culture, the blocking efficiency of wildtype C34 dropped to the level of 653+A and 653+Q (about 60-70% inhibition). Moreover, mutant C34 (647+A) added after 30 min showed similar level of inhibition as added immediately after co-culture (Fig.3B), suggesting less efficient and slower association of mutant C34 compared to the wild type peptide. At both time points, 647+A showed less efficient inhibition than 653+A and 653+Q. This finding is consistent with the hypothesis that zipping between CHR and NHR begins at the N-terminal region of CHR, and 647+A (with more upstream mismatch) blocked fusion less efficiently. The data also suggested that mutant C34 peptides (653+A or 653+Q) were able to bind to NHR, although less efficient than the wild type.

**Figure3.**
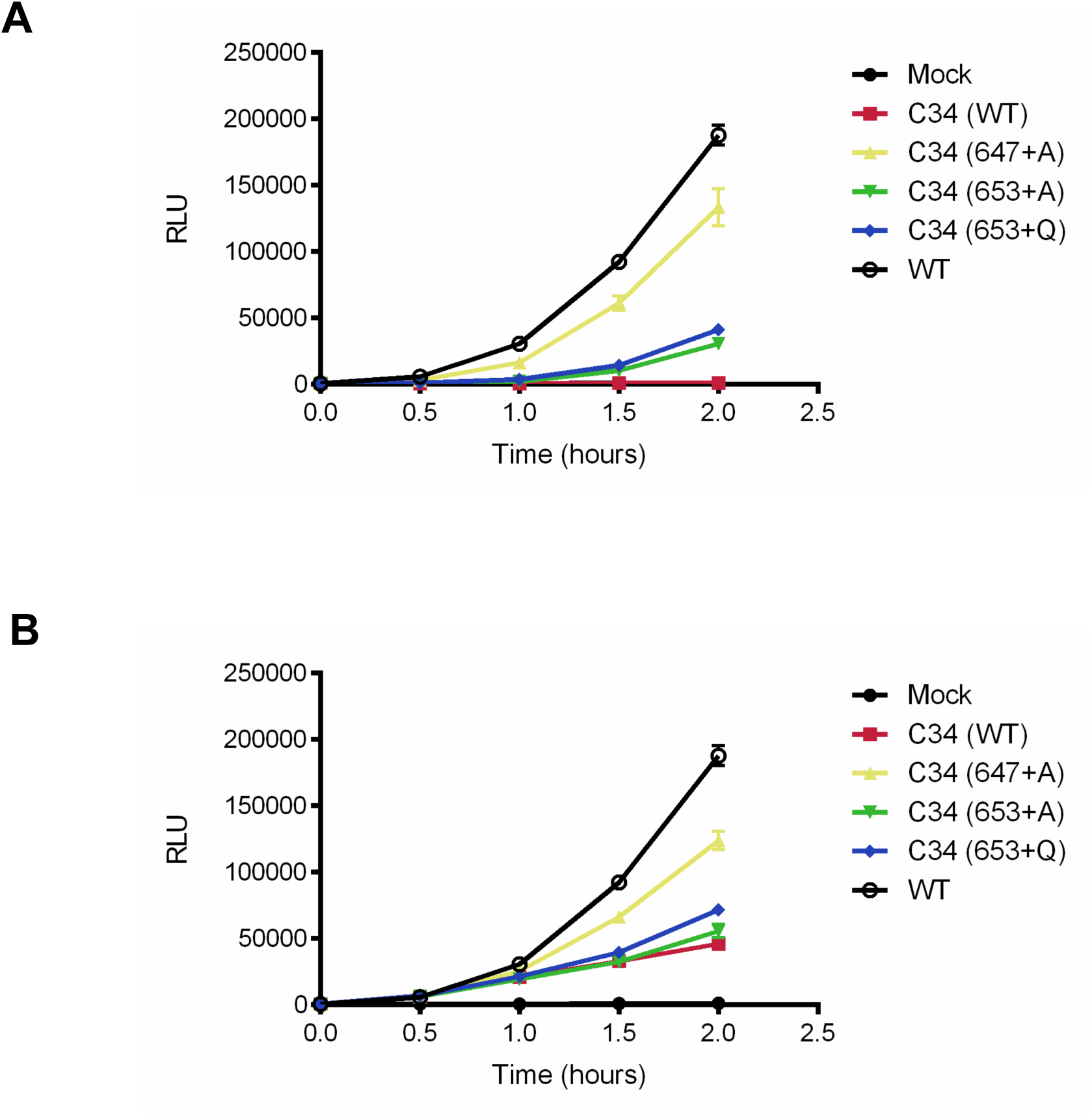
Inhibition of HIV-1 Env-mediated membrane fusion by C34 or C34-equivalent mutant peptides (647+A, 653+A, or 653+Q) evaluated by DSP assay. Expression vectors for Env were transfected into effector cells expressing DSP1–7 (293FT/DSP1–7). After approximately 24h, Env-expressing effector cells were resuspended and mixed with the target cells expressing DSP8–11(293CD4/DSP8–11). Peptides (4µM) were added at the beginning of co-culture (A) or 30 min after co-culture (B). Progression of membrane fusion from 0 to 2 h after co-culture was monitored every 0.5 h by DSP assay. The error bars indicate the standard deviation of three measurements.

### Analysis of longer NHR- and CHR-derived peptides

The inhibition assays demonstrated binding of the mutant C34 peptides to NHR, which may explain the uninterpretable CD profile due to inappropriate lengths of the used peptides. Therefore, we examined the CD profile of longer peptides pairs between C42 (aa 628-669) and N46 (aa 536-581), which showed different CD profiles than short peptides. The data suggested that the mutant peptides, even in their longer form, may have poor helix formation ability relative to wild type (Fig.2C). Next, we evaluated the thermal stability of the mixtures by CD at 222 nm (Fig. 2D), which showed that wild type N46/C42 was highly stable against thermal denaturation, with an apparent Tm of 96°C. Although the 653+Q mutant demonstrated similar cell-cell fusion activity as the wild type protein, its Tm value was much lower than wild type (59.8°C). Furthermore, the Tm of 653+A was 56.8°C, only slightly lower than 653+Q, while the 653+A mutant showed no fusion activity. Taken together, these results showed that the Tm values were not well correlated with the membrane fusion activities of the mutants.

### Effect of substitutions of Q with other amino acid residues in 653+A

We examined whether other amino acid residue (E, R, I, and L) insertion could rescue the defective fusion phenotype of 653+A. Analysis of the protein profiles of these substitutions showed similar profiles to wild type (Fig.4A and B). Mutants with E and R insertions did not improve the fusion activity at all. By contrast, substitution with I and L recovered fusion activities detected by the syncytia formation and DSP assays (Fig.5A and B). The data suggested that the amino acid residues with charged side chains were not able to rescue fusion activity. Moreover, the polar side chain that is able to form a hydrogen bond at position 653 did not appear to be essential for the recovery of membrane fusion in the 653+X context.

**Figure 4.**
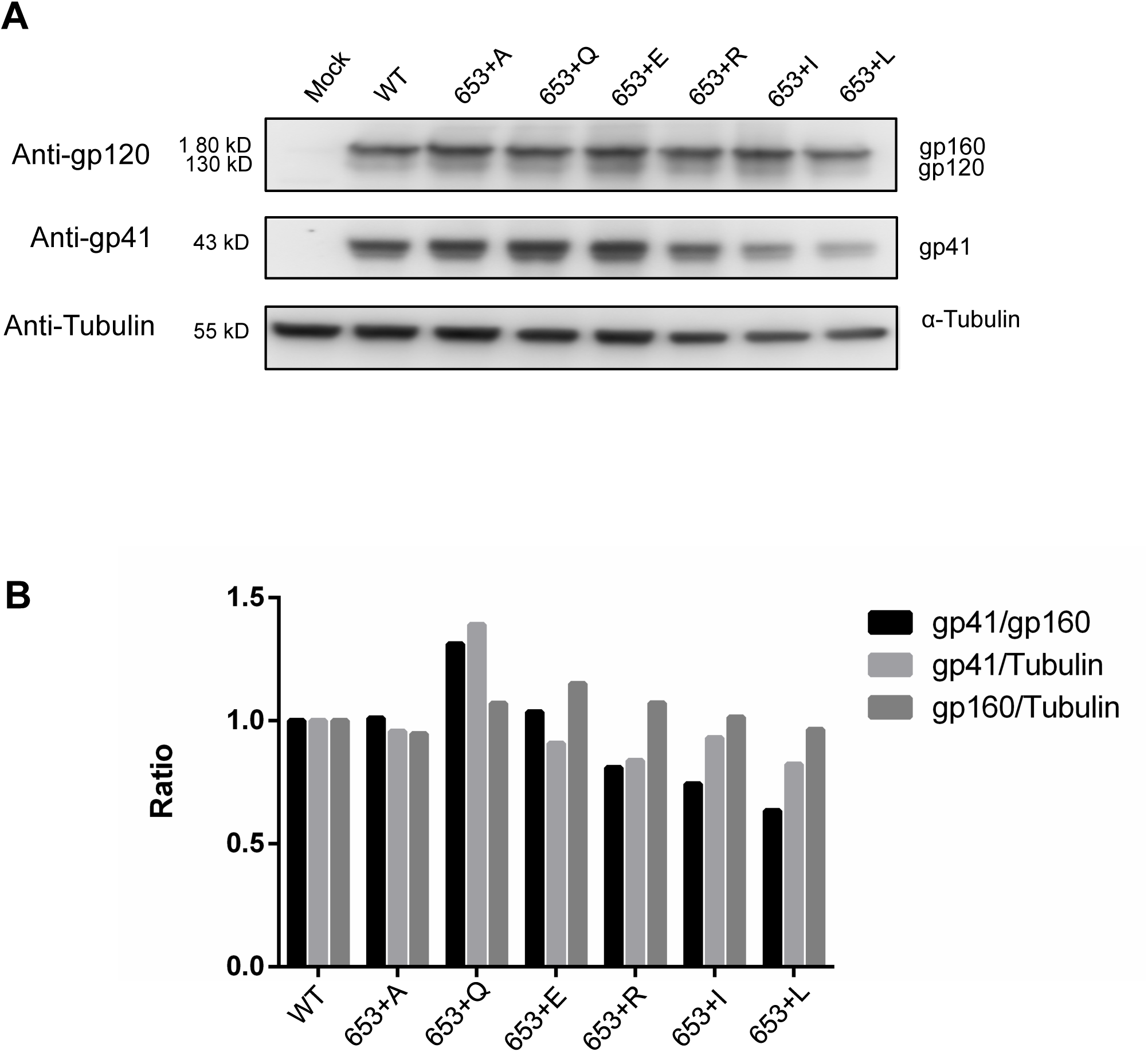
Protein profiles of Env mutants. The expression and processing of Env were analyzed by immunoblotting. (A)Representative immunoblot of Env with 653+X (X: different amino acid residue) mutants expressed in the transfected 293FT cells. Blots probed with anti-gp120 or anti-gp41 antibodies are shown.(B)The gp160, gp41, and control α-tubulin bands were quantified using Image J software. The ratios of gp41/gp160 (processing of Env) or gp160 orgp41/Tubulin (expression level of Env) were determined.

**Figure 5.**
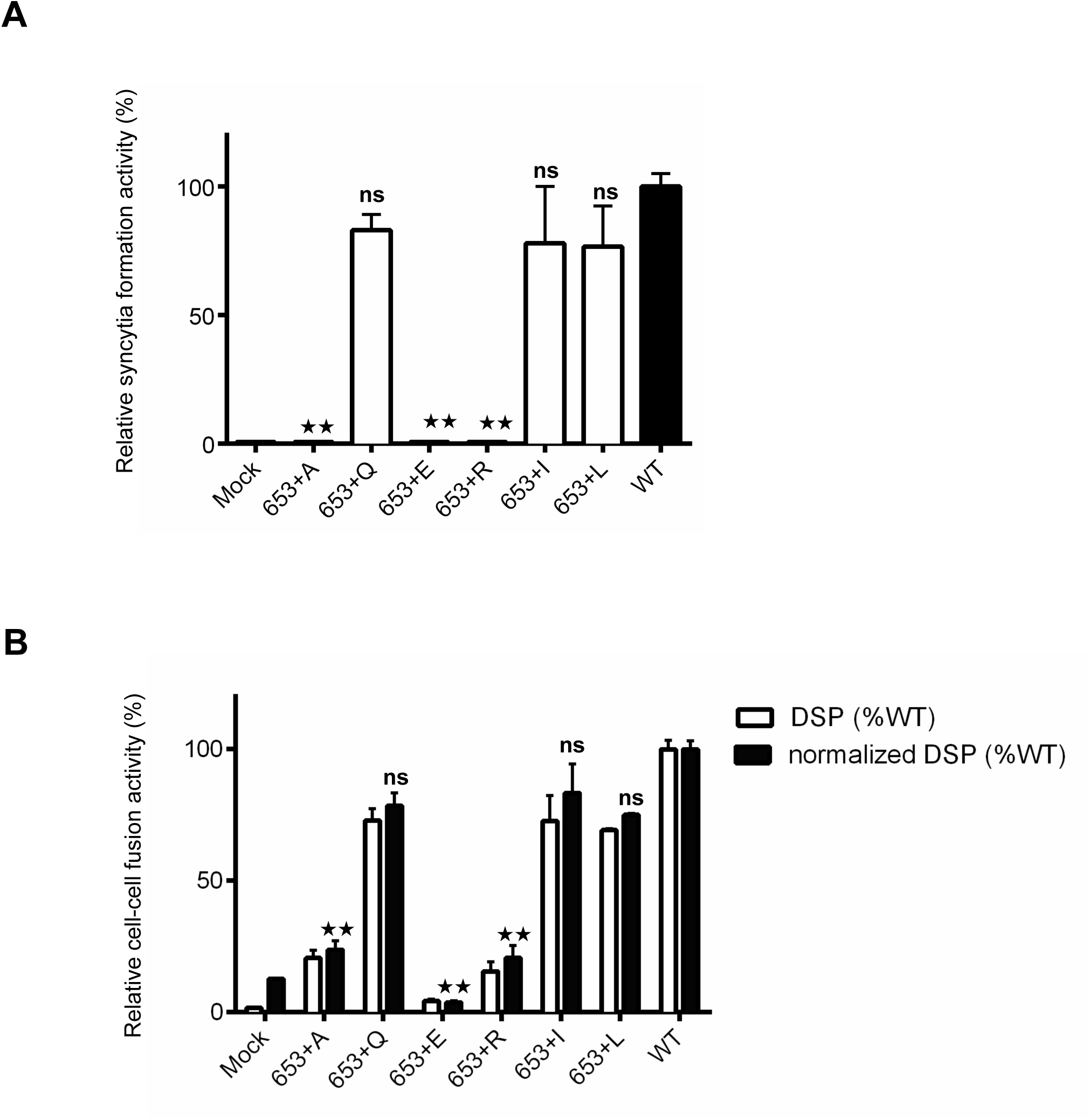
Effect of different amino acid insertion at position 653 on fusion activities. (A) Syncytia formation assay of the mutants. The relative degree of syncytia formation was shown by dividing the number of nuclei in the syncytia by the total number of nuclei in the observed fields. The wild type value was set as 100%. Error bars represent the standard deviation of results from three fields. Statistical significance was calculated by t test.**: p<0.01; ns indicates no significant difference vs. wild-type (WT) protein. (B) DSP assay to detect fusion pore formation by the mutants. The raw DSP values (white bars) and normalized values by the surface level of Env (black bars) are shown. The relative fusion activity based on RL activity is shown as white bars. **: p< 0.01, ns: no significant difference. The error bars indicate the standard deviation of triplicate experiments.

## Discussion

In this study, insertion mutations, 653+A and 653+Q, in the distal portion of HXB2gp41CHR were created. As expected from the high level of conservation of the amino acid sequences in this region, the phenotype was identical to the corresponding mutation in JRFL gp41, in which 653+A showed a severe fusion defect, while 653+Q showed recovery of activity (22).

To explore the difference between 653+A and 653+Q mutants, corresponding synthetic CHR peptides were used. Although N36 and C34 have been used widely to analyze 6HB formation (27-29), we were unable to obtain interpretable CD data using these peptides. Because our mutation is an alanine insertion rather than alanine substitution, this may have resulted in a more destabilizing effect on 6HB formation with shorter peptides. The use of N46 and C42 improved the outcome. However, the biophysical data of 653+A and 653+Q peptides in CD profiles and thermal stability did not explain the significant difference in the mutant fusion profiles. Interestingly, thermal stability of 653+Q was much lower than wild type, although both showed similar fusion activity. Furthermore, the difference in Tm between 653+Q and 653+A was only a few degrees. These data suggest that thermal stability of the complex formed by NHR and CHR region peptides may not be a good indicator of the fusion phenotype. Alternatively, under the fusion assay temperature (37°C), both mutants (653+A and 653+Q) and wild type 6HB may show comparable stability. Therefore, the difference in fusion activity between 653+A and 653+Q may be driven by the kinetics of 6HB formation rather than the stability of the formed 6HB.We cannot rule out the possibility that the use of peptides, even in the longer form, still exhibited limitations and was unable to predict the fusion profile of a native Env.

Synthetic C34 peptide is expected to competitively bind with its endogenous target, NHR within gp41, in a dominant-negative manner to inhibit 6HB formation (30-32). In our *in vitro* inhibition assay, the mutant C34 (653+A and 653+Q) showed similar weak inhibition compared to wild type C34, suggesting that the N-terminal portion of C34 may be more critical for inhibition than its distal portion. Based on the current hypothesis that 6HB formation may proceed from regions close to the connecting loop between NHR and CHR, amino acid residues at position 653 of the distal portion of CHR may not have a significant impact on the inhibitory activity of C34.

Although our mutation was generated by artificial insertional mutagenesis, comparison of the different substitution mutants may provide some useful information on the structure-function relationship of 6HB.While the side chain backbones of R and Q are quite similar, the outcome was quite different, suggesting that positive charge was not favorable in our construct, and a negative charge was worse. The presence of hydrophobic γ-carbon (absent in A, but present in Q, I, L, and R) may have been beneficial for the functional 6HB in our mutants. However, even in the presence of γ-carbon, the negative charge at this position was not favorable as mentioned above. In a previous study, we speculated that the clustering of Q, a polar and hydrophilic amino acid that can form a complex hydrogen-bond network in the 6HB, may be important for membrane fusion (33-36). However, our results of insertion mutants suggested that non-polar hydrophobic residues such as I and L were able to rescue the fusion defect, which does not support this hypothesis. Thus, the significance of the clustering of Q residues and hydrogen bond network in membrane fusion remains elusive. To obtain further insight, structural analyses of these mutants should be conducted in the future.

## Methods

### Plasmid construction

The HXB2 envelope gene sequence used in this study was codon-optimized for mammalian cell expression. A QuickChange Site-Directed Mutagenesis kit (Stratagene, La Jolla, CA, USA) was used to generate the mutants in this study. Briefly, the plasmid pHIV-env-opt HXB2was used as a template. Complementary oligonucleotide pairs containing an insertion codon were used. The mutant identified by sequencing was cloned back into the pHIV-env-opt HXB2 as the *BsrGI* and *NotI* fragment. After cloning, the entire *BsrGI*-*NotI* region, including the vector junction, was verified by sequencing. Polymerase chain reaction (PCR) was performed using Pfu Turbo (Stratagene).

### Cell culture and transfection

In brief, 293FT cells (Invitrogen, Carlsbad, CA, USA) or 293CD4 cells (293 cells constitutively expressing human CD4) were grown in Dulbecco’s modified Eagle’s medium (DMEM, Sigma, St. Louis, MO, USA) supplemented with 10% fetal bovine serum (FBS, Hyclone Labs, Logan, UT, USA). Cells were kept under 5% CO_2_in a humidified incubator (Sanyo, Kyoto, Japan) and transferred to 6- or 96-wellplatesone day before transfection. Further, Fugene HD (Fugene HD [μL]:DNA [μg]:DMEM [μL] = 5:2:200;Promega, Madison, WI, USA) was used for transient transfection. The transfection mix was incubated for 15 min at room temperature prior to addition to the cell culture in a drop-wise manner (10 μL/well). After a certain transfection time, the transfected cells were subjected to further analyses, as indicated below.

### DSP assay

To quantify cell-cell fusion, the DSP assay was performed, as described previously (15,16).Briefly, 293FT (1.5 × 10^4^) cells expressing DSP_1–7_ (293FT/DSP_1–7_) were plated in 96-well plates (Perkin Elmer Life Sciences, Waltham, MA, USA) with DMEM supplemented with 10% FBS. Subsequently, 293FT/DSP_1–7_cells were transfected with expression vectors of interest in triplicate to generate effector cells. To prepare the target cells, 293CD4/DSP_8–11_cells, a stable cell line expressing CD4 and DSP_8–11_, were seeded in 6-cm dishes (BD Falcon, San Jose, CA, USA) with the required amount of cells (cell number equal to the number of wells multiplied by 1.5 × 10^4^).The effector and target cells were mixed 20 h post-transfection at 37°C in fresh medium containing 60μMmembrane-permeable substrate for *Renilla* luciferase (RL) (Enduren Live Cell Substrate; Promega)1 h prior to mixing. RL activity was measured at the indicated times using a GloMax-Multi Plus Detection System (Promega). The effect of the peptide on membrane fusion activity was evaluated by the DSP assay. The peptide to be tested was added (final concentration 4μM) at the indicated time after coculture of the effector and target cells (1.5×10^4^ cells each). Further, the effects of the peptide were estimated by measuring RL activity at 2h after coculture.

### Syncytia formation assay

Fusion activity was further evaluated by a syncytia formation assay. Expression vectors for Env were transfected into 293CD4 cells using FuGENE HD. Unless otherwise indicated, visible syncytia formation was evaluated at 16–24 h post-transfection. To visualize nuclei, Hoechst 33342 (0.2 mg/mL; Invitrogen) was applied at 37°C for 15 min. After labeling, the cells were washed three times with 200 μL of prewarmed DMEM/FBS, and images were captured using a Fluorescent Inverted microscope (Olympus IX71). The number of nuclei in syncytia was counted to estimate the degree of syncytia formation.

### Peptide synthesis and CD analysis

The following peptides were synthesized:

N36, SGIVQQQNNLLRAIEAQQHLLQLTVWGIKQLQARIL;

C34, WMEWDREINNYTSLIHSLIEESQNQQEKNEQELL;

C34 (647+A), WMEWDREINNYTSLIHSLIAEESQNQQEKNEQELL;

C34 (653+A), WMEWDREINNYTSLIHSLIEESQNQAQEKNEQELL;

C34(653+Q), WMEWDREINNYTSLIHSLIEESQNQQQEKNEQELL;

N46,TLTVQARQLLSGIVQQQNNLLRAIEAQQHLLQLTVWGIKQLQARIL;

C42, WMEWDREINNYTSLIHSLIEESQNQQEKNEQELLELDKWASL;

C42(653+A),WMEWDREINNYTSLIHSLIEESQNQAQEKNEQELLELDKWASL;

C42(653+Q), WMEWDREINNYTSLIHSLIEESQNQQQEKNEQELLELDKWASL

The peptides were synthesized by GL Biochem Limited (Shanghai, China) with a purity of more than 95%. Protein concentrations were determined by measuring the absorbance at 280 nm in the same buffer.CD spectra were acquired on a Chirascan Plus CD spectrometer (Applied Photophysics Limited, Surry, UK) with a thermoelectric sample temperature controller. Samples for wavelength spectra were prepared as 10 μM peptide in phosphate-buffered saline (PBS). The cuvette had a 0.1 cm path length. The wavelength dependence of molar ellipticity, [θ], was monitored as the average of three scans, using a 2.5-s integration time at 1.0-nm wavelength increments. Spectra were baseline-corrected against the value for the cuvette with buffer alone. Thermal stability was determined in the same buffer by measuring [θ]_222_ as a function of temperature. A cell with a path length of 0.1cm was used with continuous stirring. Thermal melts were monitored in 2°C intervals with a 2-min equilibration at the desired temperature and an integration time of 30 s. The midpoint of the thermal unfolding transition (melting temperature, *Tm*) of each complex was determined from the maximum of the first derivative, with respect to the reciprocal of the temperature, of the [θ]_222_values(37).

### Immunoblotting

In brief, 293FT cells (2 × 10^5^) were transiently transfected with the Env expression vector using FuGENE HD in a 6-well culture plate. Cells were lysed with 60 μL of RIPA lysis buffer (Thermo Fisher Scientific, Waltham, MA, USA) at 30 h after transfection. After centrifugation at 20,000 × *g* for 30 min at 4°C, the supernatant was collected, separated by sodium dodecyl sulfate polyacrylamide gel electrophoresis on 5– 20% gels (Bio-Rad Ready Gel J), and transferred to polyvinylidene fluoride membranes (Immobilon-PSQ; Millipore, Billerica, MA, USA). The blots were probed with goat anti-gp120 polyclonal antibodies (Fitzgerald, Concord, MA, USA), the monoclonal antibody Chessie 8 (a gift from Dr. Gregory Melikyan, Department of Pediatrics, Infectious Diseases, Emory University), or anti α-Tubulin monoclonal antibody (Wako, Osaka, Japan). Donkey anti-goat IgG-horseradish peroxidase (HRP; Santa Cruz Biotechnology, Santa Cruz, CA, USA) or goat anti-mouse IgG-HRP (Santa Cruz Biotechnology) was used as the secondary antibody. The blot was further treated with an ECL Western Blot Kit (CWBIO, Beijing, China). Images were obtained using the LAS3000 system (Fujifilm, Tokyo, Japan). Band intensities were analyzed using ImageJ software (National Institutes of Health, Bethesda, MD, USA).

### CELISA

In brief, 293FT cells (2 × 10^4^ cells) were plated in 96-well View Plates (Perkin Elmer Life Sciences, Waltham, MA, USA). Five hours later, cells were cotransfected with an Env expression plasmid of interest and RL-DSP_1–7_ expression plasmids. After 40 h of transfection, cells were fixed in PBS with 4% paraformaldehyde at 25°C for 5 min and treated with 3% hydrogen peroxide in PBS at 25°C for 10 min to deactivate endogenous peroxidases. Fixed cells were incubated with 2% ECL Prime Blocking Reagent (GE Healthcare, Piscataway, NJ, USA) in PBS at 25°C for 30 min. After blocking, cells were incubated with a saturating amount of 2G12 (13.1 μg/mL, 1:1000 dilution) at 25°C for 1 h and then incubated with anti-human IgG-HRP at 1:5000 dilution (Santa Cruz Biotechnology) at 25°C for 1 h. Antibodies were diluted in 0.2% ECL Prime Blocking Reagent. Luminescence was detected using ECL Prime Western Blotting Detection Reagent (GE Healthcare) and a GloMax 96 microplate luminometer (Promega). Before calculation of the relative expression levels, the average background signal from blank control wells was subtracted from the blank control well.

### Statistical analysis

Statistical analyses were performed with Prism software (GraphPad, La Jolla, CA, USA). Unless otherwise specified, differences among groups were assessed using *t* test and were considered significant at *p*<0.01.

## Acknowledgement

This work was supported in part by the Research Program on Emerging and Re-emerging Infectious Diseases and the Program of Japan Initiative for Global Research Network on Infectious Diseases from the Japan Agency for Medical Research and Development under Grant JP19fm0108006h0005.

